# A novel, dichloromethane-fermenting bacterium in the *Peptococcaceae* family, ‘*Candidatus* Formamonas warabiya’, gen. nov. sp. nov.

**DOI:** 10.1101/2020.08.09.233494

**Authors:** Sophie I Holland, Haluk Ertan, Michael J Manefield, Matthew Lee

## Abstract

Dichloromethane (DCM; CH_2_Cl_2_) is a toxic groundwater pollutant that also has a detrimental effect on atmospheric ozone levels. As a dense non-aqueous phase liquid, DCM migrates vertically through groundwater to low redox zones, yet information on anaerobic microbial DCM transformation remains scarce due to a lack of cultured organisms. We report here the characterisation of strain DCMF, the dominant organism in an anaerobic enrichment culture (DFE) that is capable of fermenting DCM to the environmentally benign product acetate. Stable carbon isotope experiments demonstrated that the organism assimilated carbon from DCM and bicarbonate via the Wood-Ljungdahl pathway. Strain DCMF is the first anaerobic DCM-degrading bacterium also shown to metabolise non-chlorinated substrates. It appears to be a methylotroph utilising the Wood-Ljungdahl pathway for metabolism of methyl groups from methanol, choline, and glycine betaine, which has implications for the flux of climate-active compounds from subsurface environments. Community profiling and enrichment of the cohabiting taxa in culture DFE to the exclusion of strain DCMF suggest that it is the sole organism in this culture responsible for substrate metabolism, while the cohabitants persist via necromass recycling. Genomic and physiological evidence support placement of strain DCMF in a novel genus, ‘*Candidatus* Formamonas warabiya’.

## Introduction

Dichloromethane (DCM, CH_2_Cl_2_) is one of the most commonly encountered subsurface pollutants in industrial areas (1). Current global production of DCM exceeds 900 Gg y^-1^, of which 70% is manufactured by humans (2). The remaining 30% comes from natural sources including biomass burning, oceanic sources, and geothermal activity (2). Due to widespread production and use of DCM, both surface and tropospheric levels of this toxic chemical continue to rise (3–6). Atmospherically, DCM has recently been recognised as a potent greenhouse gas with detrimental effects on ozone (7). The compound also poses a threat to human health (8,9) and microbial function (10–12).

Microbial transformation of DCM is an option for remediation of oxic and anoxic environments. Aerobic DCM transformation is catalysed by a DCM dehalogenase found in facultative methylotrophs (13,14), but the enzyme responsible for anaerobic DCM dechlorination has not yet been identified. DCM is a dense non-aqueous phase liquid that descends through groundwater to low redox zones, and so anaerobic degradation plays a vital role in its removal from contaminated sites. Yet study of anaerobic DCM degradation has been hindered by the highly limited number of organisms capable of this metabolism. *Dehalobacterium formicoaceticum* strain DMC is the only isolate (15) and from the handful of enrichment cultures (16–18), only ‘*Candidatus* Dichloromethanomonas elyunquensis’ strain RM has been characterised (19,20). Both *D. formicoaceticum* and ‘*Ca*. Dichloromethanomonas elyunquensis’ are obligate anaerobic DCM-degrading bacteria and have genome sequences available (15,19,21,22). A combination of genomic, physiological and proteomic work has demonstrated the central role of the Wood-Ljungdahl pathway (WLP) in DCM metabolism in both organisms, however variations on the pathway result in different end products (15,20,23). *D. formicoaceticum* ferments DCM to formate and acetate in a 2:1 molar ratio (15), whilst ‘*Ca*. Dichloromethanomonas elyunquensis’ completely mineralises DCM to H_2_ and CO_2_ (23).

A new anaerobic DCM degrading bacterium, strain DCMF, was recently subjected to whole genome sequencing and also encoded a complete WLP (24). Strain DCMF is the dominant organism in a non-methanogenic bacterial consortium, designated culture DFE (24). The community was enriched from an organochlorine-contaminated aquifer near Botany Bay, Sydney, Australia and culture DFE has been maintained on DCM as the sole energy source (24). However, genome-based metabolic modelling suggested that strain DCMF may have a wider substrate repertoire due to the presence of 81 full-length MttB superfamily methyltransferases and glycine/betaine/sarcosine reductase genes (24).

Here, we report that strain DCMF is the first non-obligate anaerobic DCM degrading bacterium and characterise its metabolism of DCM, quaternary amines, and methanol, whilst also considering the role of the cohabiting bacteria in culture DFE. Stable carbon isotope labelling was used to determine the fate of DCM carbon and function of the WLP. Based on its genomic and physiological novelty, strain DCMF is proposed to form a novel genus within the *Peptococcaceae* family. Using contemporary molecular and traditional cultivation techniques (including exclusion cultivation – removal the dominant taxon), this study represents a thorough and robust characterisation of a novel bacterium despite its presence in a multi-lineage enrichment culture. This supports recent initiatives to redefine how bacterial lineages are formally recognised (25).

## Materials and Methods

### Culture medium

Culture DFE was grown in anaerobic, defined bicarbonate-buffered mineral salts medium as previously described (24). To investigate the requirement for exogenous bicarbonate during DCM degradation, cultures were instead buffered with 3-morpholinopropane-1-sulfonic acid (MOPS, 4.2 g l^-1^), either with or without 4 mM NaHCO_3_. To study the metabolic fate of DCM, ^13^C-labelled DCM ([^13^C]DCM, 1 mM) was used. To study the assimilation of inorganic carbon, ^13^C-labelled bicarbonate (NaH^13^CO_3_, 5 mM) was added to MOPS-buffered culture medium.

To test alternative growth substrates, DCM was replaced with the following (5 mM unless stated otherwise): carbon monoxide (2 mM), choline chloride, dibromomethane, dimethylglycine, formic acid, H_2_, glycine betaine, methanol, sarcosine, syringic acid, trimethylamine. Cultures amended with choline chloride, glycine betaine, and trimethylamine were also amended with the following compounds as electron acceptors (15 mM unless otherwise stated): fumarate (80 mM, tested with trimethylamine only), NaNO_2_, NaNO_3_, Na_2_SO_3_ and Na_2_SO_4_. Acetate, H_2_, and lactate were tested as electron donors with Na_2_SO_3_ and Na_2_SO_4_ as electron acceptors. Glycine betaine and sarcosine (5 mM) were tested as electron donors with H_2_ (10 mM) as electron acceptor.

### Analytical methods

DCM, dibromomethane, acetate, formate, methanol, and trimethylamine were quantified using a Shimadzu Plus GC-2010 gas chromatograph with flame ionisation detector (GC-FID) equipped with a headspace autosampler (PAL LHS2-xt-Shim; Shimadzu, Rydalmere, Australia; Table S1). HCO_3_^-^(as gaseous CO_2_) and H_2_ were quantified using a Shimadzu GC-2010 gas chromatograph with pulsed discharge detector (GC-PDD; Table S1). In all analyses, the inlet temperature was 250°C, split ratio 1:10, FID temperature 250°C or PDD temperature 150°C.

Choline and glycine betaine were quantified using liquid chromatography with tandem mass spectrometry (LC-MS/MS). The Agilent 1200 Series LC (Agilent Technologies, Mulgrave, Australia) was fitted with a Luna C18(2) column (150 × 4.6 mm, 5 µm; Phenomenex, Lane Cove West, Australia). The mobile phases were 0.5 mM ammonium acetate in water (A) and 100% methanol (B). Samples (5 µl) were eluted with a linear gradient from 95:5 (A:B) to 0:100 (A:B) over 10 min, then held at 0:100 (A:B) for 1 min. The LC was coupled to an Applied Biosystems QTRAP 4000 quadrupole mass spectrometer (SCIEX, Mulgrave, Australia) and electrospray ionization performed in the positive mode. The machine was operated in multiple reaction monitoring (MRM) mode and the following precursor-product ion transitions were used for quantification: *m/z* 104.0 → 59.0 (choline) and *m/z* 118.0 → 57.7 (glycine betaine).

Labelled and unlabelled acetate, CO_2_, and HCO_3_^-^ were quantified via GC with triple quadrupole mass spectrometry (GC-TQMS) performed with an Agilent 7890A GC system (Table S1). The TQMS was operated in MRM mode identifying the following precursor-product ion transitions: *m/z* 43 → 15.2 (unlabelled acetate), *m/z* 44 → 15.1 ([1-^13^C]acetate), *m/z* 44 → 16 ([2-^13^C]acetate), *m/z* 45 → 16.1 ([1,2-^13^C]acetate), *m/z* 45 → 29 (^13^CO_2_), *m/z* 44 → 28 (^12^CO_2_).

GC-TQMS in MRM mode was also used to quantify dimethylamine, methylamine, sarcosine, and glycine, using alanine as an internal standard. Following derivatisation (26) (Table S1), the following precursor-product ion transitions were used: *m/z* 117.2 → 89.1 (dimethylamine), *m/z* 103.2 → 74.9 (methylamine), *m/z* 116.2 → 44.1 (sarcosine and alanine), and 102 → 30.1 (glycine).

### Bacterial quantification

Genomic DNA was extracted from 2 ml liquid culture as previously described (24). Strain DCMF and total bacterial 16S rRNA genes were quantified via quantitative real-time PCR (qPCR) with primers Dcm775/Dcm930 and Eub1048/Eub1194 (27), respectively (Table S2). Standard curves were prepared by making serial 10-fold dilutions of plasmid DNA carrying cloned strain DCMF 16S rDNA or *Dehalococcoides* sp. 16S rDNA (for total bacterial quantification). Reactions were carried out on a CFX96 thermal cycler (Bio-Rad) and the data was analysed with CFX Maestro v1.0 software (Bio-Rad). Strain DCMF 16S rRNA gene copy numbers were converted to cell numbers by dividing by four (the number of 16S rRNA genes in the genome).

### 16S rRNA gene amplicon sequencing

Community profiling was carried out on the above DNA samples. The 16S rRNA gene was amplified with the 515F/806R primer pair with adapters (Table S2). Samples were sequenced with Illumina MiSeq technology by The Hawkesbury Institute for the Environment Next Generation Sequencing Facility. Amplicon reads were processed in QIIME2 (28) using the dada2 pipeline (29): forward and reverse reads were trimmed and joined, chimeras were removed, and samples were rarefied to the lowest sequencing depth. Taxonomy was assigned to genus level using a Naïve Bayes classifier trained on a full-length 16S rRNA gene SILVA database (release 133) and the lowest 1% abundant reads were filtered out. Alpha diversity was assessed with Shannon’s diversity index and pairwise comparisons made with a Kruskal-Wallis test. A two-dimensional PCA plot was created from the weighted Unifrac distance matrix. Samples were compared by the proportion of substrate consumed, as well as timepoint, to account for differing metabolic rates between substrates (Table S3).

### Exclusion cultivation of DFE cohabitant bacteria

To eliminate strain DCMF and enrich the cohabiting bacteria in culture DFE, two rounds of dilution to extinction cultures (20 ml) were set up in 30 ml glass serum bottles (Fig S1). These were prepared with the standard medium amended with one of: casamino acids (5 g l^-1^), ethanol (10 mM), glucose (10 mM), peptone (5 g l^-1^), 1-propanol (10 mM), yeast extract (5 g l^-1^). Following qPCR confirmation that strain DCMF was below the limit of detection in the lowest active dilution culture, these cultures were subject to Illumina 16 rRNA gene amplicon sequencing and used to inoculate triplicate microcosms amended with one of: 1 mM DCM, 5 mM choline chloride, or 5 mM glycine betaine (Fig S1), which were monitored for eight weeks.

### Fluorescence in situ hybridisation microscopy

Fluorescence in situ hybridisation (FISH) was carried out with a strain DCMF-specific oligonucleotide probe (Dcm623, 5’-/Cy3/CTCAAGTGCCATCTCCGA-3’), designed using ARB (30), and probe Eub338i (5’-/6-FAM/GCTGCCTCCCGTAGGAGT-3’) (31) to target all bacteria. FISH was carried out as per an established protocol for fixation on a polycarbonate membrane, using minimal volumes of reagents (32). Cells were fixed with protocols for both Gram negative (31) and Gram positive cell walls (33). Hybridisation was carried out with a formamide-free buffer. Cells were counterstained with VECTASHIELD® Antifade Mounting Medium containing 1.5 µg ml^- 1^ 4,6-diamidino-2-phenylindole (Vector Laboratories, Burlingame, CA, USA). Images were captured on a BX61 microscope equipped with a DP80 camera (Olympus Australia, Notting Hill, Australia) using Olympus cellSens Dimension software v2.1. Strain DCMF cell length and width was determined from a sample of 20 cells using the linear measurement tool within the program.

## Results

### Dichloromethane fermentation

Culture DFE consumed 1.9 ± 0.0 mM DCM within 35 days, yielding 3.7 ± 2.2 × 10^8^ strain DCMF cells ml^-1^ (Fig 1A), or 2.0 ± 1.2 × 10^14^ strain DCMF cells per mole of substrate consumed. The product of DCM fermentation was acetate (1.4 ± 0.1 mM; Fig 1A), which was not observed in abiotic controls. DCM attenuation did not proceed in MOPS-buffered cultures free of bicarbonate Fig 1B). However, in analogous cultures amended with 4 mM NaHCO_3_, DCM attenuation was observed, yet HCO_3_^-^ concentrations did not significantly change (p = 0.11, two-tailed t-test between days 0 and 65; Fig 1B).

**Figure 1.**
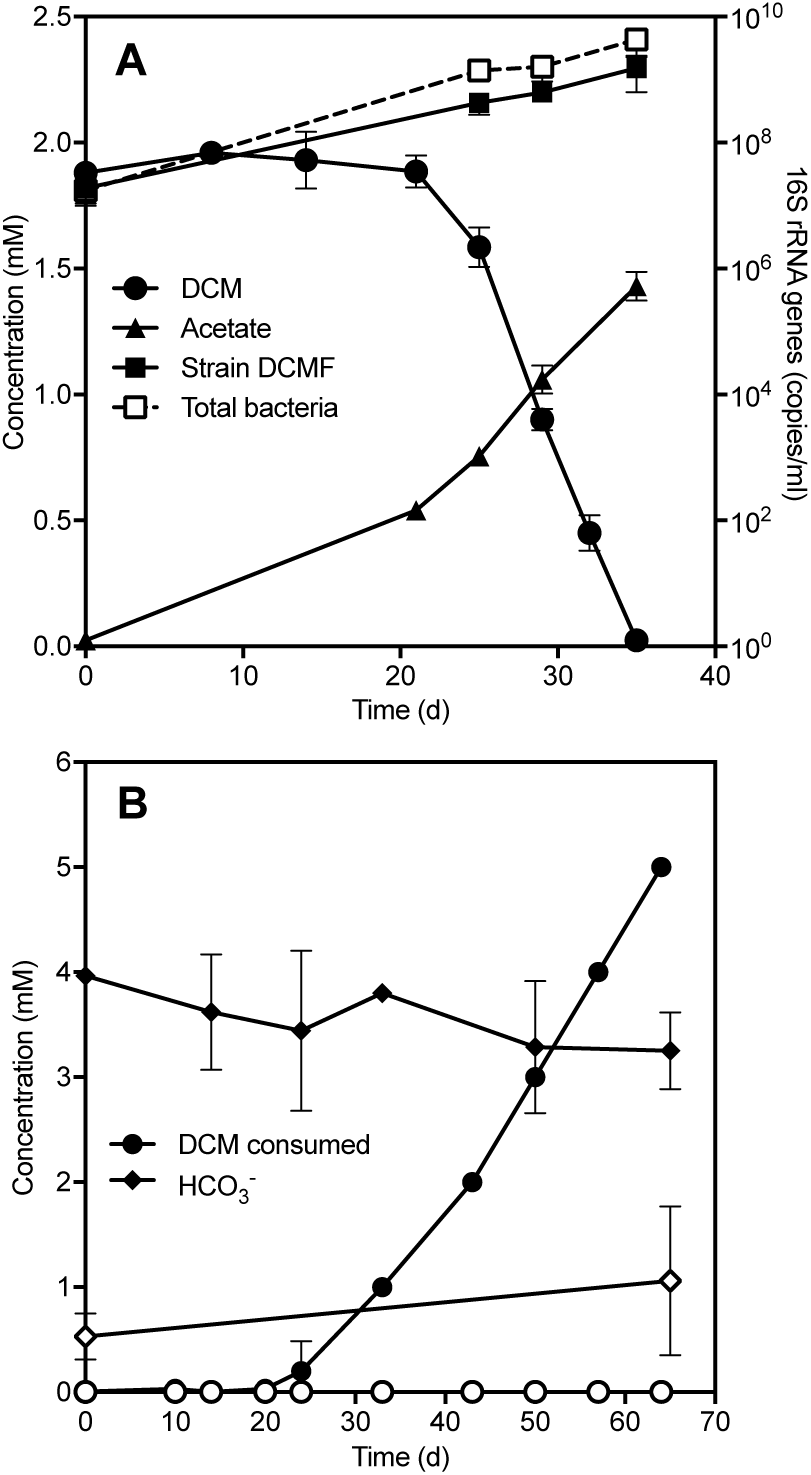
Strain DCMF ferments DCM to acetate, reliant on exogenous bicarbonate. (A) Strain DCMF growth was concomitant with the depletion of DCM and formation of acetate. Error bars represent standard deviation, *n* = 2. Substrate and product concentrations are quantified on the left y-axis; strain DCMF and total bacterial 16S rRNA gene copies are quantified on the right y-axis. (B) In MOPS-buffered medium, DCM consumption was only observed in the presence of bicarbonate (filled circles). Cumulative DCM consumption is from repeat amendment of 1 mM DCM. Empty circles represent cultures with no exogenous bicarbonate. Error bars represent standard deviation, *n* = 3.

### Metabolism of quaternary amines and methanol

Of the additional substrates tested as sole energy source or with an electron acceptor, strain DCMF growth was observed when methanol, choline or glycine betaine (5 mM each) were supplied (Fig 2). Culture DFE consumed methanol (4.3 ± 0.2 mM) over 30 days, yielding 3.1 ± 0.1 mM acetate and 2.4 ± 0.6 × 10^9^ strain DCMF cells ml^-1^ (Fig 2A). This corresponded to 5.7 ± 1.4 × 10^14^ strain DCMF cells per mole substrate utilised. No methanol depletion was observed in the abiotic (cell-free) control.

**Figure 2.**
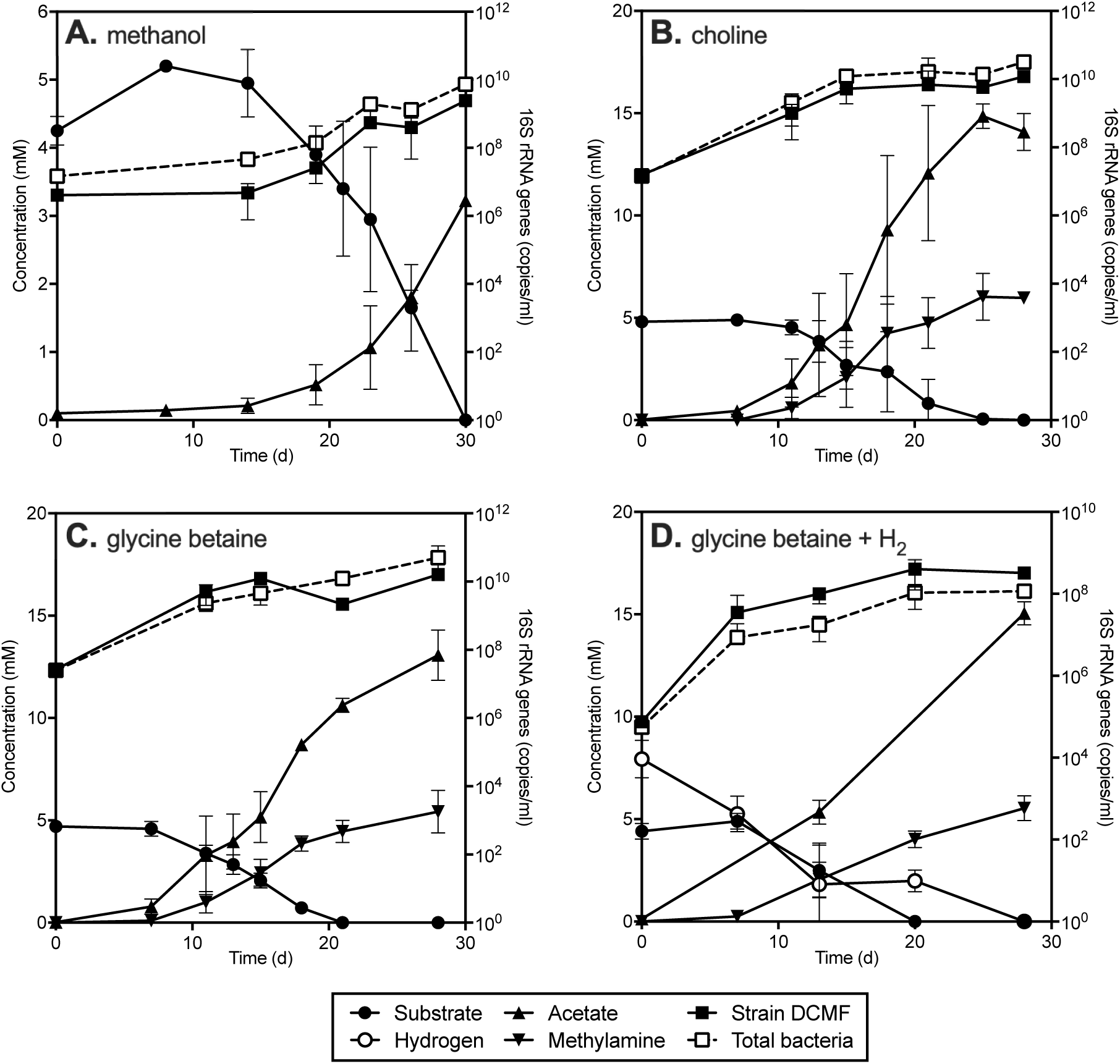
Strain DCMF growth was also correlated with the metabolism of methanol and quaternary amines. Strain DCMF growth correlated with the depletion of methanol and formation of acetate (A) and the depletion of choline (B) and glycine betaine (C) with formation of acetate and methylamine. Cultures amended with glycine betaine and hydrogen (D) did not produce trimethylamine, rather acetate and methylamine were once again the products. Substrate and product concentrations are quantified on the left y-axis; strain DCMF and total bacterial 16S rRNA gene copies are quantified on the right y-axis. Error bars represent standard deviation, *n* = 3.

Culture DFE consumed choline (4.8 ± 0.2 mM) within 25 days, producing 15 ± 0.6 mM acetate and 6.0 ± 1.1 mM methylamine (Fig 2B). Glycine betaine (4.7 ± 0.3 mM) was consumed within 21 days, with production of 11 ± 0.4 mM of acetate and 4.5 ± 0.6 mM methylamine (Fig 2C). Trimethylamine, dimethylamine, sarcosine (methylglycine), and glycine were not detectable throughout. Neither acetate nor methylamine were detected in abiotic controls, and the latter was also absent from cultures amended with DCM. Strain DCMF cell proliferation aligned with the consumption of these two substrates, yielding an increase of 1.4 ± 0.4 × 10^9^ and 5.3 ± 0.4 × 10^8^ cells ml^-1^ in choline- and glycine betaine-amended cultures, respectively (Fig 2B, C). This corresponded to 3.0 ± 0.9 × 10^14^ cells per mole of choline, and 1.1 ± 0.1 × 10^14^ cells per mole of glycine betaine utilised.

DFE cultures amended with quaternary amine metabolic pathway intermediates dimethylglycine and sarcosine + H_2_ also demonstrated production of acetate and methylamine, which again aligned with strain DCMF cell proliferation (Fig S2). Sarcosine was not degraded in the absence of H_2_ (data not shown). Following the observation of strain DCMF growth and methylamine production in cultures amended with sarcosine + H_2_, DFE cultures were also set up with glycine betaine + H_2_ to determine whether glycine betaine could be reductively cleaved to trimethylamine and acetate. These cultures consumed all glycine betaine (4.4 ± 0.4 mM) and hydrogen (7.9 ± 0.9 mM) within 28 days, producing 15 ± 0.6 mM acetate and 5.5 ± 0.6 mM methylamine, but no trimethylamine (Fig 2D). Strain DCMF cell yields (4.0 ± 2.8 × 10^8^ cells ml^-1^) were similar to that when glycine betaine was the sole energy source.

Culture DFE was unable to utilise CO, dibromomethane, ethanol, formate, syringic acid or trimethylamine as sole energy sources (no growth and/or acetogenesis observed). Strain DCMF was further unable to use any of the tested pairs of electron donors (acetate, choline, glycine betaine, H_2_, lactate, trimethylamine) and acceptors (CO_2_, fumarate, Na_2_SO_4_, Na_2_SO_3_, NaNO_2_, and NaNO_3_).

### Shifts in DFE community composition in response to substrate consumption

Community profiling with Illumina 16S rRNA gene amplicon sequencing showed that culture DFE is composed of a limited number of taxa – only 12 amplicon sequencing variants (ASVs) were present at ≥ 2% relative abundance in > 1 sample (Fig 3). Community composition was similar in cultures amended with DCM, choline, and glycine betaine, which had a common, DCM-amended inoculum (Fig 3A,B,C), but was simplified in cultures that had been maintained on methanol for two sub-cultivations and had a methanol-amended inoculum (Fig 3D; Fig S3A). While strain DCMF was the dominant organism at the time of inoculation and during substrate consumption, its relative abundance decreased markedly in the lag phase prior to substrate consumption, falling to as little as 0.96% in a methanol-amended replicate at day 14 (Fig 3). Taxa such as *Synergistaceae* (except in methanol-amended cultures, where this taxon was absent), *Desulfovibrio* and *Veillonellaceae* increased in relative abundance during this lag phase, while *Spirochaetaceae* and *Lentimicrobiaceae* increased towards the end of and following substrate depletion, particularly in quaternary amine-amended cultures (Fig 3).

**Figure 3.**
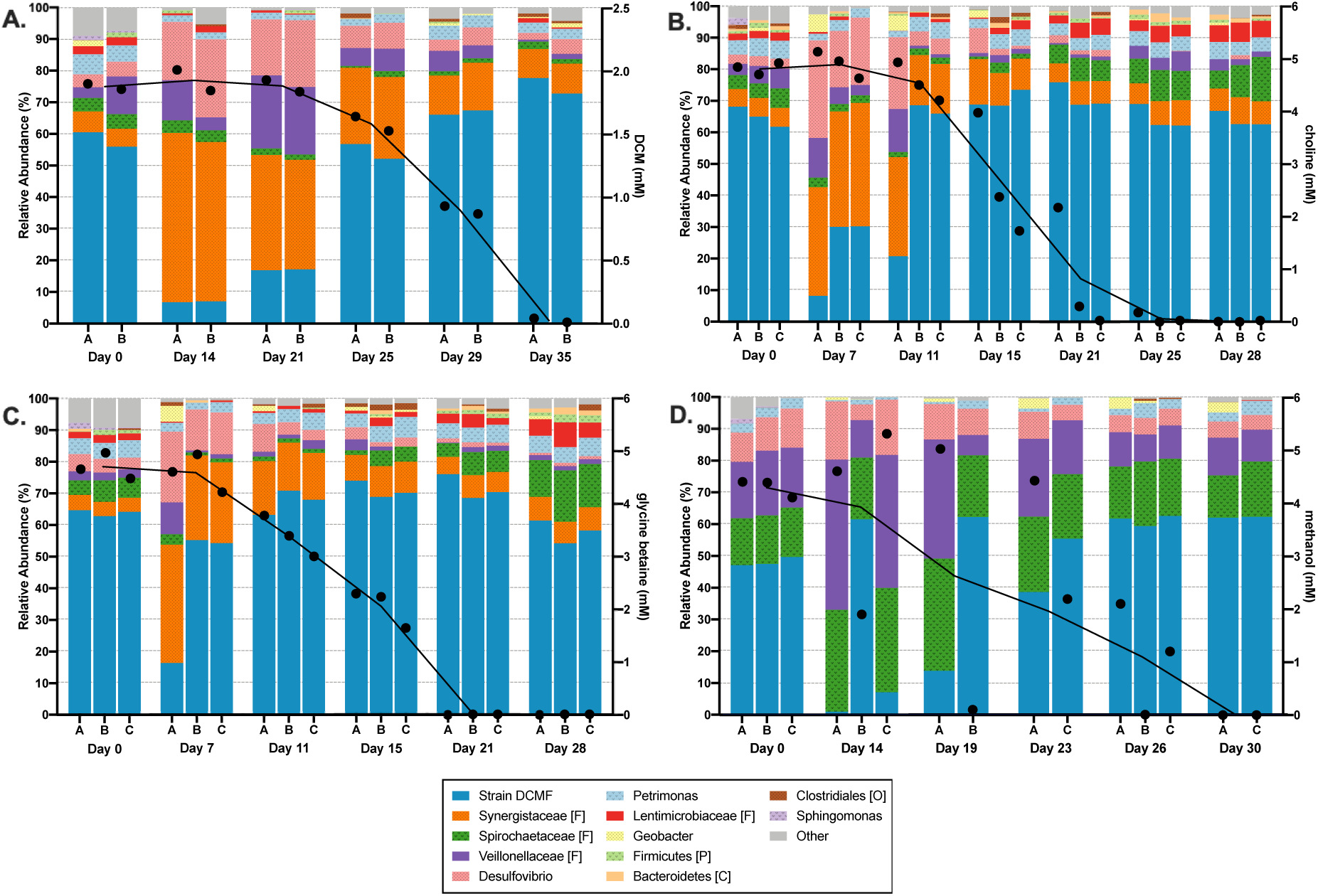
Culture DFE is subject to temporal shifts in community composition, with strain DCMF dominant during substrate degradation. Illumina 16S rRNA amplicon sequencing was used to determine DFE community composition (left y axis) at timepoints across the growth experiments amended with (A) DCM, (B) choline, (C) glycine betaine, and (D) methanol reported in Figs 1 and 2. Taxa are reported down to genus level where possible, otherwise taxonomic level is indicated in the legend ([F] = family, [P] = phylum, [C] = class, [O] = order). Reads with <1% abundance were filtered out in QIIME2. Unassigned reads and taxa consistently <2% relative abundance were classed together as ‘Other’. Substrate concentration (black circles, right y-axis) and a line connecting the mean substrate concentration at each time point is overlaid on the community composition graphs. These are aligned with the time points written on the x-axis, not drawn to scale.

Differences in the DFE community were driven by the degree of substrate consumption (defined in Table S3), more than the various substrates (Fig S3B). While there was no significant difference in the Shannon diversity index between the samples when grouped by substrate (Kruskal-Wallis p-value 0.0976; Fig S3C), there was a highly significant difference between all groups when clustered by substrate consumption (Kruskal-Wallis p-value <0.00001; Fig S3D).

### Exclusion of cohabitants as DCM and quaternary amine consumers

Attempts to isolate strain DCMF proved unsuccessful (24). Therefore, to test the hypothesis that strain DCMF was the sole consumer of DCM and quaternary amines, the cohabiting bacteria in culture DFE were enriched to the exclusion of strain DCMF (exclusion cultivation). This was achieved by dilution to extinction cultures on rich media amended with casamino acids, glucose, peptone or yeast extract. These growth conditions variously enriched *Bacillus, Desulfovibrio, Geobacter, Petrimonas*, and *Veillonellaceae*, but not strain DCMF (Fig S4E-H). *Spirochaetaceae* and *Synergistaceae* phylotypes did not grow on the tested rich media.

The strain DCMF-free cohabitant cultures were then tested for their ability to utilise DCM, choline, and glycine betaine. There was no significant substrate depletion in these cultures (Fig S4A-D), and therefore no evidence of DCM, choline, or glycine betaine degradation by the *Bacillus, Desulfovibrio, Geobacter, Petrimonas*, or *Veillonellaceae* phylotypes in culture DFE.

### Strain DCMF morphology

FISH microscopy enabled selective visualisation of strain DCMF cells, which appeared rod-shaped and occurred singly or in chains (Fig 4A). On average, strain DCMF cells were 1.69 ± 0.27 µm long and 0.64 ± 0.12 µm wide. FISH images confirmed that strain DCMF numerically dominated culture DFE during DCM dechlorination, congruent with community profiling results (Fig 4C).

**Figure 4.**
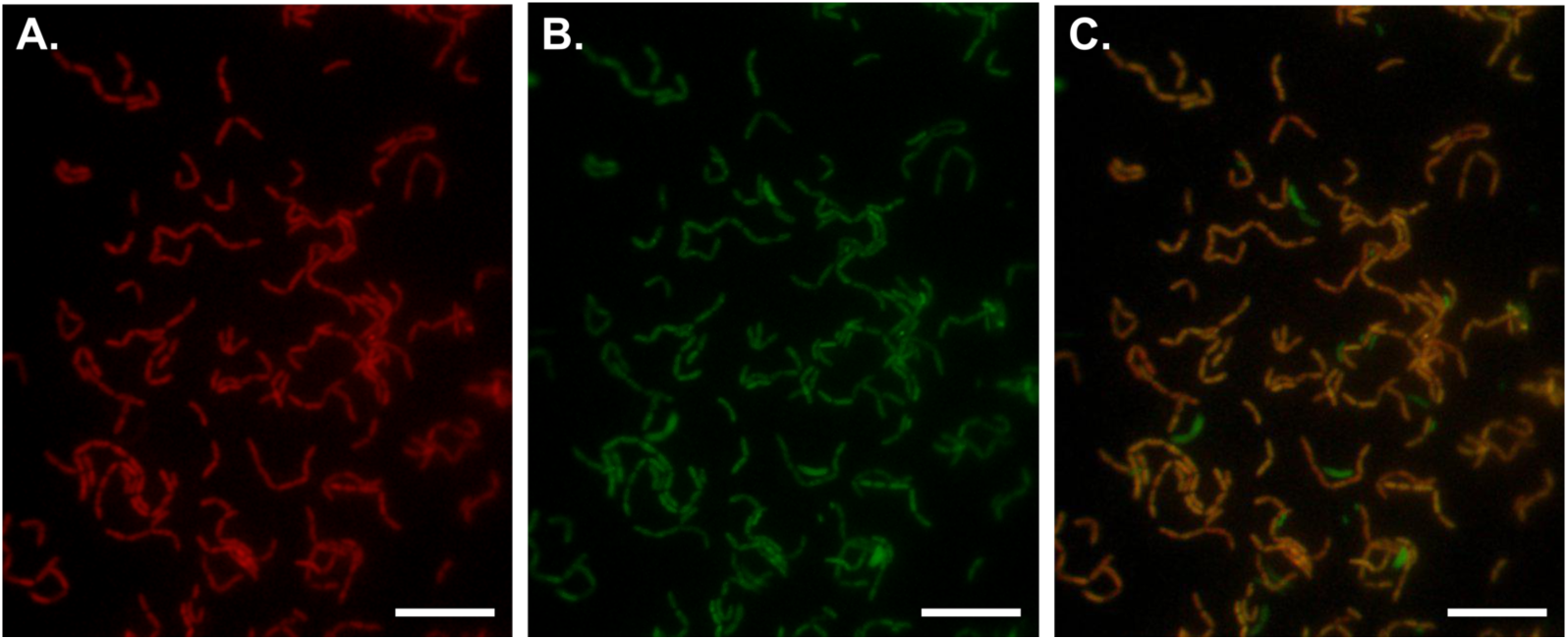
Morphology of strain DCMF. Fluorescence in situ hybridisation (FISH) microscopy images with strain DCMF cells stained red with the Cy3-labelled Dcm623 probe (A), all bacterial cells stained green with the 6-FAM-labelled Eub338i probe (B), and the overlay of Cy3- and 6-FAM-labelling in these images (C). The scale bars represent 10 µM.

### Strain DCMF is mixotrophic

To ascertain the fate of DCM carbon, triplicate DFE cultures were amended with [^13^C]DCM. When 2 700 ± 328 µM DCM had been consumed, 666 ± 160 µM of acetate was produced (Fig 5A), of which 47.1 ± 5.5% was unlabelled, 30.4 ± 2.8% was methyl group labelled ([2-^13^C]acetate), and 22.5 ± 4.3% was both methyl and carboxyl group labelled ([1,2-^13^C]acetate; Fig 5C). A ^13^C mass balance was achieved by summing the measured concentrations of ^13^C-labelled carbon in acetate (670 ± 289 µM) and H^13^CO_3^-^_ (815 ± 120 µM) with the calculated concentrations of ^13^CO_2_ in the flask headspace (982 ± 144 µM) and [^13^C]acetate equivalents in biomass (994 ± 121 µM; Fig 5B, Table S4). This amounted to 128 ± 8.2% recovery of the labelled carbon, indicating no unknown fate of DCM in culture DFE.

**Figure 5.**
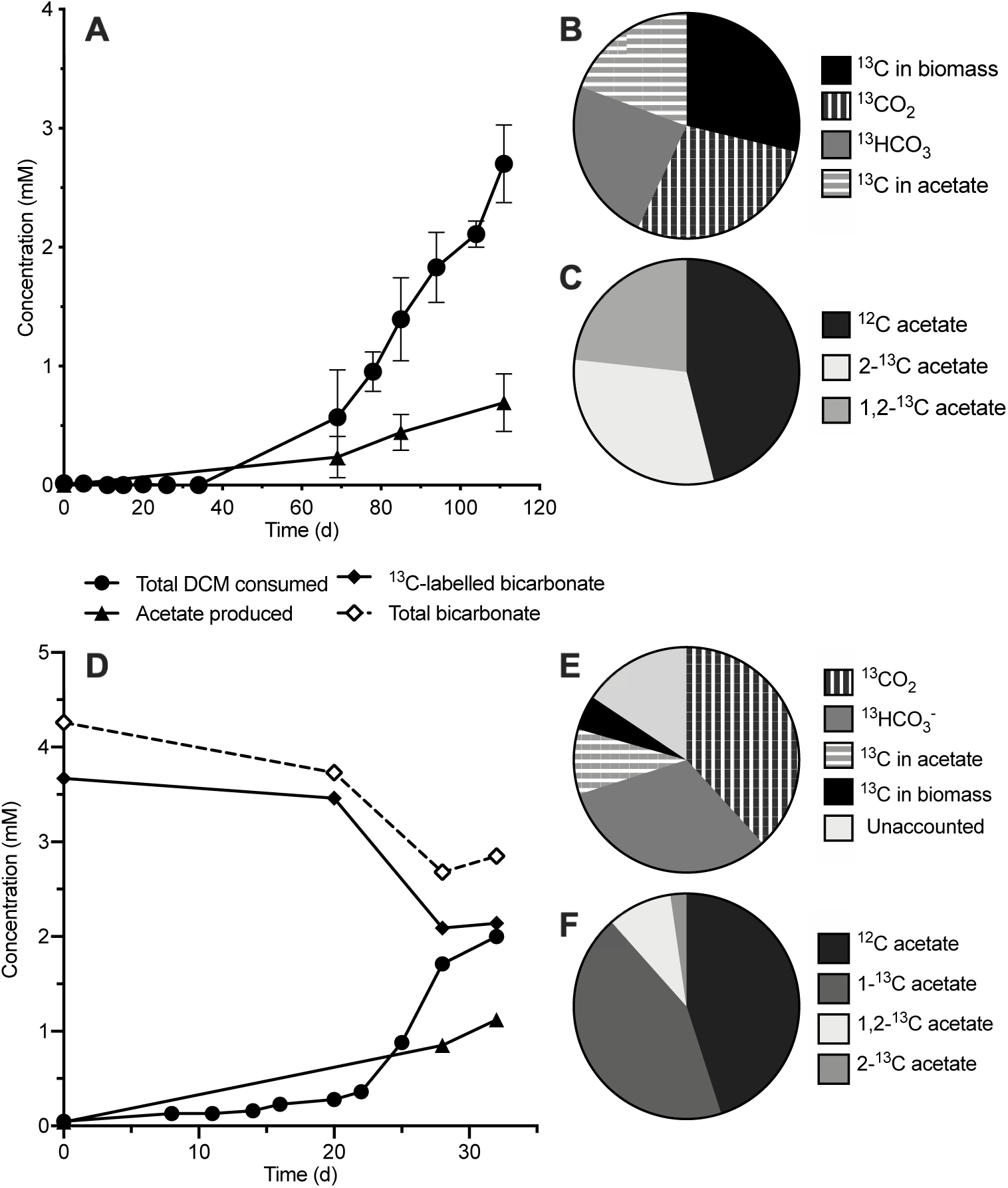
Strain DCMF assimilates carbon from DCM and bicarbonate to form acetate. (A) Cumulative [^13^C]DCM consumption with concomitant with acetate production, including the ^13^C mass balance from [^13^C]DCM (B) and proportion of labelled and unlabelled acetate (C). Error bars represent standard deviation, *n* = 3. (D) Cumulative DCM consumption and acetate production in cultures amended with H^13^CO_3_^-^. Total (labelled and unlabelled) aqueous HCO_3_^-^ is also shown (i.e. gaseous CO_2_ is not accounted for here). Values in (D) are from a single representative culture as all triplicates had similar dechlorination rates and product concentrations but began dechlorinating at different times. The ^13^C mass balance from H^13^CO_3_^-^ (E) and proportion of labelled and unlabelled acetate (F) is again shown. All pie charts represent the average of triplicate cultures at the final time point.

Analogous work was then carried out with unlabelled DCM in MOPS-buffered medium amended with ^13^C-labelled bicarbonate, showing that strain DCMF incorporated carbon from CO_2_ into the carboxyl group of acetate. The culture consumed 2000 µM DCM and 2150 ± 492 µM ^13^C from bicarbonate. It produced 973 ± 140 µM acetate (Fig 5D), of which 45.0 ± 2.3% was unlabelled, 43.5 ± 1.8% was labelled on the carboxyl group ([1-^13^C]acetate), 2.2 ± 1.3% was labelled on the methyl group, and 9.3 ± 0.1% was labelled on both carbons (Fig 5F). A mass balance indicated 84.5 ± 7.0% recovery of the labelled carbon in acetate (600 ± 84.9 µM), the remaining H^13^CO_3_^-^ (2280 ± 170 µM) and ^13^CO_2_ (2740 ± 204 µM), and strain DCMF biomass (710 ± 9.74 µM; Fig 5E, Table S4).

## Discussion

### The DFE community

Strain DCMF is a novel bacterium present in enrichment culture DFE, which has been maintained with DCM as sole external source of energy for five years and at least 20 consecutive transfers (24). Of the five other phylotypes previously reported in culture DFE, based on 16S rRNA genes identified from genome sequencing data (24), four remained amongst the most abundant in the present work (*Desulfovibrio, Lentimicrobiaceae, Spirochaetaceae* and *Synergistaceae*), while one was no longer detected (*Ignavibacteria*). In combination with the similar community profiles observed across four different substrates, this suggests that culture DFE is a long-term stable-state community.

Illumina amplicon sequencing, FISH microscopy and qPCR all supported the previous observation (24) of strain DCMF as the dominant organism in culture DFE during substrate consumption, and linked growth of strain DCMF to depletion of DCM, methanol, choline, and glycine betaine. Attempts to generate an axenic culture of strain DCMF have been unsuccessful, similar to the DCM-mineralising bacterium ‘*Ca*. Dichloromethanomonas elyunquensis’ in culture RM (19,23). How the cohabiting organisms in both cultures persist despite numerous transfers and addition of only a simple chlorinated compound (DCM) to minimal, anaerobic medium is a question of interest. While hydrogenotrophic acetogens and methanogens form major sub-populations in culture RM (18,19,23), culture DFE is non-methanogenic (24) and was unable to grow on H_2_+CO_2_ alone.

Instead, the timing of the changes in relative abundance and known substrate range of major phylotypes in culture DFE suggest that the cohabiting bacteria use cellular detritus resulting from expired strain DCMF cells as an energy source (i.e. necromass fermentation). Some of the most abundant cohabiting phylotypes in culture DFE – *Desulfovibrio, Bacteroidetes* (containing the families *Lentimicrobiaceae* and *Petrimonas*), *Spirochaetes/Treponematales, Synergistetes* – have previously been associated with hydrocarbon and organohalide-degrading mixed cultures (34– 38), although their abundance was not linked to degradation of the primary substrate (with the exception of some *Desulfovibrio* species) and some reports also suggested that they persist via necromass recycling (16,37–39).

Five of the 12 taxa in culture DFE were categorically excluded from being primary metabolisers of DCM, choline, and glycine betaine when tested in the absence of strain DCMF (Fig S4). *Spirochaetaceae* and *Synergistaceae* phylotypes could not be enriched to the exclusion of strain DCMF. However, their relative abundance during growth on DCM, choline, and glycine betaine diminished relative to strain DCMF, suggesting that it is unlikely that they are primary consumers of these substrates. This needs to be confirmed by proteomic assessment of the DFE community.

### The role of the WLP in DCM metabolism

Amongst anaerobic DCM-dechlorinating bacteria, strain DCMF is unique in producing solely acetate as an end product (Fig 1A). *D. formicoaceticum* produced formate and acetate in a 2:1 molar ratio (15), while ‘*Ca*. Dichloromethanomonas elyunquensis’ completely mineralised DCM to H_2_, CO_2_ and Cl^-^ (23). The latter organism is unique in also encoding and expressing reductive dehalogenases during growth with DCM (19,20). Despite these differences, both organisms utilise the WLP for DCM metabolism (15,20,23) as is likely the case with strain DCMF. Removal of bicarbonate from the culture medium precluded DCM dechlorination and ensuing work with ^13^C- labelled DCM and bicarbonate demonstrated that strain DCMF is mixotrophic, i.e. assimilates carbon from both DCM and CO_2_, similar to *D. formicoaceticum* (23).

These experiments also provided compelling evidence for the transformation of DCM to a WLP intermediate, mostly likely methylene-tetrahydrofolate (CH_2_=FH_2_; Eq. 1). The production of H^13^CO_3_^-^ from [^13^C]DCM suggested that CH_2_=FH_2_ is disproportionated into the WLP where it is oxidised to HCO_3_^-^ (Eq. 2, Fig 6). The electrons released then reduce the remaining CH2=FH_2_ into the methyl group of acetate (Eq. 3). However, the production of unlabelled acetate (47%) indicates that the excess unlabelled HCO_3_^-^ (30 mM) in the medium is an alternative electron acceptor to CH2=FH2 for acetogenesis (Eq. 4; Fig 6). The reduction of HCO_3_^-^ to acetate requires twice as many electrons for acetate synthesis than CH2=FH2 (i.e. eight vs. four). Taking this ratio into account, along with ∼1:1 ratio of unlabelled to labelled acetate suggests that approximately 67% of electrons derived from DCM oxidation were directed toward HCO_3_^-^ reduction and 33% to CH_2_=FH_2_.

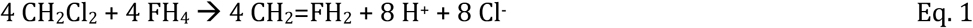

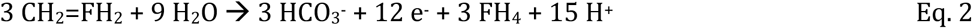

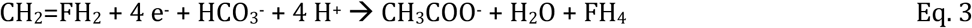

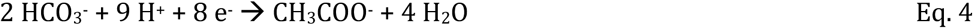

**Figure 6.**
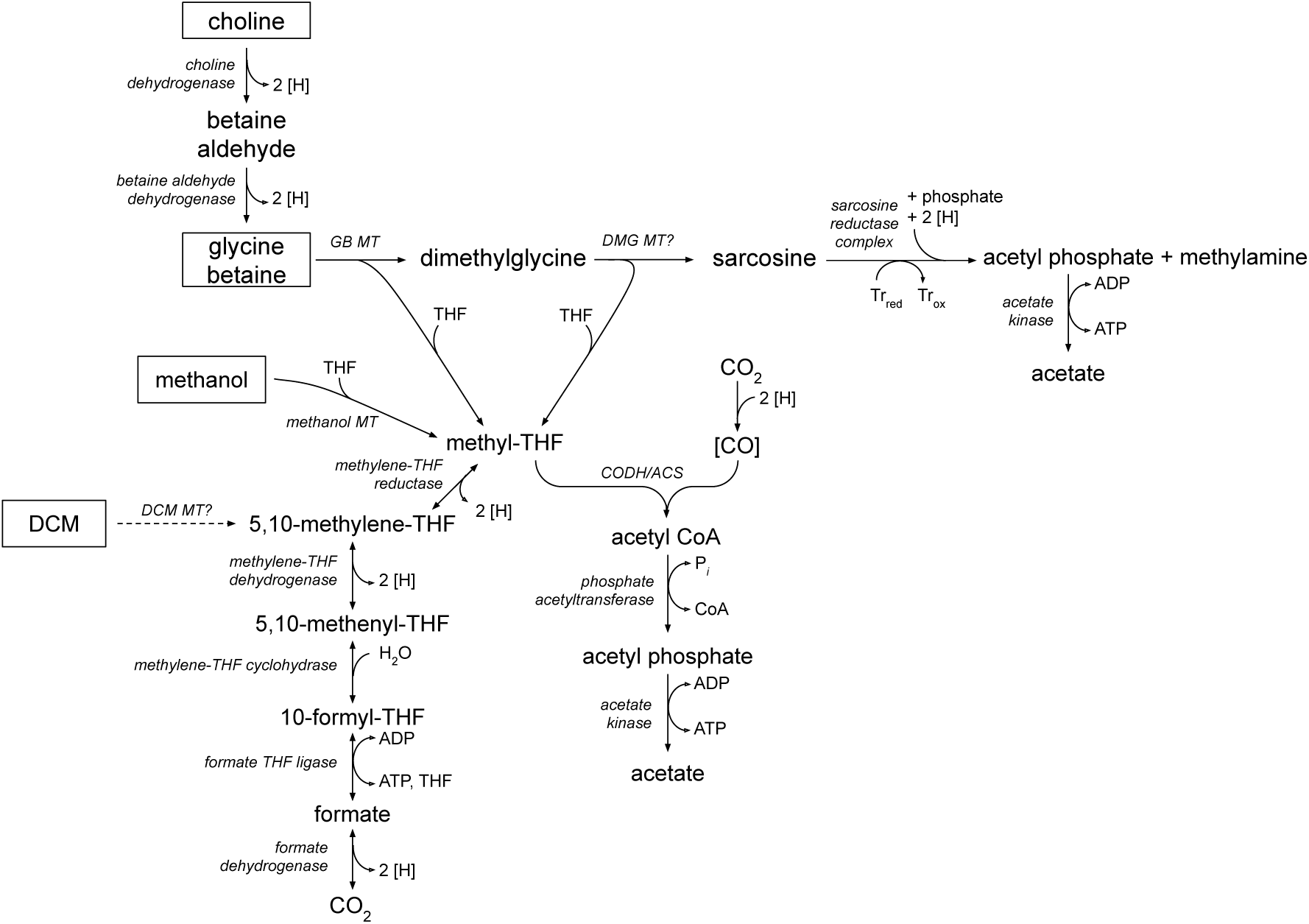
Proposed model for metabolism of DCM, methanol and quaternary amines by strain DCMF. The Wood-Ljungdahl pathway is central to transformation of all substrates into acetate. Proteins (with the exception of that catalysing the putative transformation of DCM to CH_2_=FH_2_, indicated by dotted arrow) are all identified in the strain DCMF genome and listed in Table S5. Abbreviations: CODH/ACS, carbon monoxide dehydrogenase/acetyl-CoA synthase; DCM, dichloromethane; DH, dehydrogenase; DMG, dimethylglycine; GB, glycine betaine; MT, methyltransferase; ox, oxidised; red, reduced; THF, tetrahydrofolate; Tr, thioredoxin.

The production of [1,2-^13^C]acetate from [^13^C]DCM is consistent with the reduction of H^13^CO_3_^-^ outlined above. However, the proportion (22.5%) was surprisingly high, given the relatively small contribution that labelled H^13^CO_3_^-^ from 2.7 mM [^13^C]DCM would make to the 30 mM unlabelled HCO_3_^-^ present in the culture medium. It is possible that co-localisation of WLP proteins in the cytoplasm may cause the reduction of H^13^CO_3_^-^ at a higher ratio than expected (i.e. 9%). Studies with [^13^C]DCM in *D. formicoaceticum* detected the ^13^C label solely in the methyl group of acetate ([2-^13^C] acetate), congruent with DCM oxidation stopping at formate (23,40), while studies with another *Dehalobacterium* species in mixed culture that was capable of formate oxidation similarly detected [1,2-^13^C]acetate (41).

DFE cultures amended with unlabelled DCM and ^13^C-labelled HCO_3_^-^ in MOPS-buffered medium produced an analogous proportion of [1-^13^C]acetate. A similar proportion of acetate (45.0%) to that observed in the [^13^C]DCM work was unlabelled, in this case evidently formed using unlabelled HCO_3_^-^ produced from DCM. Thus, the ^13^C-labelling experiments support the hypothesis that DCM metabolism involves the WLP and are consistent with the oxidation of formate to HCO3^-^. As an exogenous supply of formate was unable to stimulate growth of culture DFE, strain DCMF alone is likely responsible for formate metabolism, which contrasts with the inability of *D. formicoaceticum* to further transform this metabolite (15). The production of HCO_3^-^_ from formate balances with its uptake during acetogenesis, congruent with a net flux of approximately zero, leading to the proposal that DCM is transformed as per Equation 5.

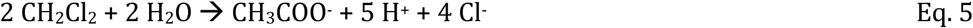

### Metabolism of non-chlorinated substrates

Strain DCMF is the first non-obligate anaerobic DCM-degrading bacterium. A genome-based metabolic model previously suggested that the abundance of MttB superfamily methyltransferases (named for their founding member, a trimethylamine:corrinoid methyltransferase) encoded by strain DCMF may permit growth on methylamines amines and/or glycines (24). While strain DCMF was unable to metabolise trimethylamine, growth was observed with glycine betaine and the closely related compound choline. Both compounds are quaternary amines with significant environmental roles. Glycine betaine is an osmoprotectant widely used by bacteria (42–44), marine algae (45), marine invertebrates (46), plants (47), and some vertebrates (48). It is also an important source of nitrogen, comprising up to 20% of the total nitrogen in hypersaline environments (49). Choline is typically more abundant, albeit as a part of larger molecules including eukaryotic phospholipids, and can be converted to glycine betaine by a near ubiquitous pathway in soil and water environments (50).

Accordingly, strain DCMF encodes both the choline dehydrogenase (Ga0180325_11215) and betaine aldehyde dehydrogenase (Ga0180325_114191) required for this transformation to glycine betaine. Based on the stoichiometry of observed end products, growth on putative pathway intermediates, and genomic information, we propose that strain DCMF likely stepwise demethylates glycine betaine to dimethylglycine and then sarcosine (methylglycine), which is then reductively cleaved to form acetate (via acetyl-phosphate) and methylamine (Supplementary Discussion and Fig 6). The electron equivalents produced from demethylation can be used for additional reduction of CO_2_ to acetate via the WLP, as well as the reductive cleavage of sarcosine. This combination of demethylation and reductive cleavage has previously only been observed in *Sporomusa* spp. (51,52) and is a novel metabolic pathway within the *Peptococcaceae* family. A theoretical energy balance of the product formation and strain DCMF cell yield suggested that no other organisms in culture DFE were involved in quaternary amine metabolism (Supplementary Discussion).

The strain DCMF genome also encodes a number of methanol methyltransferases, which are likely utilised for transformation of methanol into CH_2_=FH_4_ prior to its entry into the WLP (Supplementary Discussion and Fig 6).

### Environmental significance

The ability of strain DCMF to utilise choline, glycine betaine and methanol suggests that its environmental relevance extends beyond DCM contaminated sites. Coastal salt marshes and intertidal mudflats represent significant sources of methane from the demethylation of trimethylamine, which is in turn derived from quaternary amines (53–55). Both trimethylamine and methanol are non-competitive methane precursors, which may allow large methanogen populations to develop in environments where sulphate reduction would typically dominate (56,57). Indeed, trimethylamine is responsible for 60 - 90% of methane production in coastal salt marshes and intertidal sediments (54,56). The transformation of quaternary amines to methylamine by strain DCMF provides a pathway of lower methanogenic potential that could operate in coastal subsurface environments. Strain DMCF does create acetate as a major end product, which can be utilised by acetoclastic methanogens. However, unlike methylated amines, methanogens have to compete with more thermodynamically favourable processes such as sulphate reduction for this substrate.

Furthermore, DCM has recently also been recognised as a potent greenhouse gas with ozone-depleting potential (7), and oxygenated hydrocarbons such as methanol can influence atmospheric ozone formation through reactions with nitrous oxides (58). Therefore, although DCM, methanol, and quaternary amines are seemingly disparate substrates, they are closely linked to the atmospheric flux of climate-active gasses from anoxic, subsurface environments. This is both via the direct influence that DCM and methanol can have on ozone, and the indirect influence of quaternary amines on the flux of methylated amines and methane.

### Provisional classification of strain DCMF as a novel genus and species

In addition to previously reported 16S rRNA gene phylogeny (24), whole genome analysis of universally conserved marker genes and amino acid identity methods showed the closest relative of strain DCMF to be *D. formicoaceticum* strain DMC (Supplementary Methods and Results). While the former two methods support the placement of strain DCMF in a novel genus, the latter was on the borderline of suggested thresholds (Supplementary Results). However, the physiological information presented here distinguishes strain DCMF from the sole representative of the genus *Dehalobacterium*, which has thus far only proved capable of growth on DCM (15). Strain DCMF also harbours a significantly larger genome than *D. formicoaceticum* (6.44 Mb for the former, 3.77 Mb for the latter) (22), which may account for its wider substrate range. Strain DCMF appears to be an anaerobic methylotroph, capable of metabolising a unique range of one-carbon compounds (DCM, methanol) or substrates from which it can utilise methyl groups (choline, glycine betaine, dimethylglycine, sarcosine). Thus, multiple lines of evidence support the placement of strain DCMF within a novel genus in the family *Peptococcaceae*. As strain DCMF is not yet represented in pure culture despite intensive efforts to isolate the organism, we propose it be classified in the *Candidatus* category (59) as ‘*Candidatus* Formamonas warabiya’ strain DCMF gen. nov. sp. nov.

### Description of ‘Candidatus *Formamonas’ gen. nov*

‘*Candidatus* Formamonas [Form.a.mon’as. L. adj. *formicum* relating to formic acid or, more generally, one-carbon compounds; Gr. n. *monas* unit; ML. n. *Formamonas* the one-carbon utilising unit.

‘*Candidatus* Formamonas’ is strictly anaerobic and metabolises one-carbon and methylated compounds including DCM, methanol and quaternary amines glycine. Methylene/methyl groups are metabolised via incorporation into the WLP. The type species is ‘*Candidatus* Formamonas warabiya’.

### Description of ‘Candidatus *Formamonas warabiya’ sp. nov*

‘*Candidatus* Formamonas warabiya [war.a.bi’ya N.L. n. *warabiya* the Dharawal name for the area between Botany Bay and Bunnerong, honouring the Traditional Custodians of the land where this bacterium was sampled from]. Permission was granted from the Dharawal Language Program research group for use of this word as the species name.

Utilises DCM, methanol, choline, glycine betaine, dimethylglycine as sole sources of electrons under anoxic conditions. Can also utilise the electron donor and acceptor pair H_2_ and sarcosine. The aforementioned substrates plus CO_2_ are carbon sources. The primary fermentation product is acetate. Methylamine is also produced from choline, glycine betaine, dimethylglycine, and sarcosine + H_2_. The WLP is likely used for carbon fixation and metabolism of the methyl groups removed from substrates. Cells are rod shaped (1.69 × 0.27 μm).

Type strain DCMF^T^ is not available in pure culture. The source of inoculum was contaminated sediment from the Botany Sands aquifer, adjacent to Botany Bay, Sydney, Australia. The type material is the finished genome of ‘*Candidatus* Formamonas warabiya’ strain DCMF, which is 6.44 Mb and has a G+C content of 46.4% (GenBank accession number CP017634.1; IMG genome ID 2718217647).

## Abbreviations

DCM: dichloromethane
FISH: fluorescence *in situ* hybridisation
GC-FID: gas chromatography with flame ionisation detector
GC-PDD: gas chromatography with pulse discharge detector
GC-TQMS: gas chromatography with triple quadrupole mass spectrometry
LC-MS/MS: liquid chromatography with tandem mass spectrometry
MRM: multiple reaction monitoring
WLP: Wood-Ljungdahl pathway

## Acknowledgements

We are grateful to Kate Montgomery in the School of Biotechnology and Biomolecular Science, UNSW, for her assistance with FISH microscopy. Thanks to Dr Valentina Wong for her initial culturing assistance. SH was supported by an Australian Government Research Training Program Scholarship.

