## Supplementary Text and Figures for "A novel, dichloromethane-fermenting bacterium in the *Peptococcaceae* family, ‘*Candidatus* Formamonas warabiya’, gen. nov. sp. nov."

### Supplementary Methods

Whole genome taxonomic analysis of strain DCMF was carried out with the GTDB-Tk (Genome Taxonomy Database toolkit) (60). The average nucleotide identity (ANI) tool from the Kostas lab (61) was used to calculate ANI values between strain DCMF and *D. formicoaceticum* strain DMC. CompareM (https://github.com/dparks1134/CompareM) was used to calculate the two-way average amino acid identity (AAI) between the anaerobic DCM degraders and other related bacteria in the family *Peptococcaceae*.

### Supplementary Results

Whole genome taxonomic analysis of strain DCMF with the GTDB-Tk identified its closest relative as *D. formicoaceticum* strain DMC, placing them together in the novel family taxon *Dehalobacteriaceae* (order *Dehalobacteriales*, class *Dehalobacteriia*, phylum *Firmicutes*). The GTDB re-classified a wide range of bacterial taxa based on its analysis pipeline, including splitting the traditional class of *Clostridia* (which includes the family *Peptococcaceae*) into a variety of more specific, monophyletic classes (62), and hence this classification is equivalent to the assignation of family *Peptococcaceae* previously suggested for strain DCMF (24).

Strain DCMF had 77.19% ANI to its closest relative, *D. formicoaceticum* strain DMC. Given that ANI offers robust resolution primarily above 80% values (61), AAI analysis was instead carried out to evaluate genomic distance between strain DCMF and its closest relatives. *D.* formicoaceticum was again confirmed as the closest relative to strain DCMF (AAI value 66.54%), with '*Ca.* Dichloromethanomonas elyunquensis' and other members of the *Peptococcaceae* all considerably lower (Table S6). AAI between strain DCMF and *D. formicoaceticum* is at the lower end of the AAI genus boundary suggested by Konstantinidis and Tiedje (63), but within the more recent boundary of 55-60% proposed by Rodriguez-R and Konstantinidis (61).

### Supplementary Discussion

*Necromass fermentation by cohabiting bacteria in culture DFE*

Of the cohabiting phylotypes in culture DFE, many have been shown to ferment compounds that would facilitate necromass recycling. *Synergistaceae* are well-known amino acid fermenters (64,65) and can be saccharolytic (66). It was expected they would be enriched with casamino acids, however they appear to have been outcompeted by *Veillonellaceae*. *Spirochaetes* are commonly detected in anoxic environments contaminated with hydrocarbons and organohalides (67,68) and a recent report demonstrated their role as necromass recyclers in such ecosystems (38). It was therefore expected that the was expected that the *Synergistaceae* in culture DFE would be enriched with casamino acids, and the *Spirochaetes* with glucose, however they appear to have been outcompeted by the faster-growing *Veillonellaceae* and *Petrimonas* phylotypes, respectively (Fig S4).

Members of the *Veillonellaceae* are typically associated with animal and/or human hosts and have been reported to ferment carbohydrates and metabolise organic acids and amino acids (69). *Petrimonas* and *Lentimicrobiaceae* are both members of the phylum *Bacteroidetes* with isolates shown to ferment a range of carbohydrates (70–72).

The sulphate-respiring genus *Desulfovibrio* has a broad substrate range and has been associated with contaminated environments both as a primary degrader (35,73) and a synergistic cohabitant consuming fermentation products of other organisms (18,37,68,74). A *Desulfovibrio* species is also present in an anaerobic DCM-dechlorinating culture dominated by *Dehalobacterium*, where it likely consumes formate produced from DCM metabolism and may also be capable of reducing minor amounts of sulphate present in the yeast extract added to the medium (41). Given that culture DFE is unable to consume exogenous formate and that the *Desulfovibrio* phylotype in culture DFE was enriched to apparent purity on peptone (Fig S4C), it may persist here via a proteolytic metabolism.

*Metabolism of choline by strain DCMF*

When used as a substrate for growth, choline is typically metabolised into trimethylamine in anoxic subsurface environments (54,55,75), via a choline-trimethylamine lyase (CutC) (76). The inability of culture DFE to utilise exogenously provided trimethylamine and absence of this compound in choline-amended cultures suggested that a different metabolic pathway was being utilised here. Furthermore, the sole CutC homolog (Ga0180325_112585) in the strain DCMF genome contains only three of the six conserved residues predicted to be necessary for catalytic activity in other bacteria (76), is ~50 codons shorter than characterised CutC proteins, and not located within a bacterial microcompartment gene cluster.

Direct demethylation of choline to dimethylethanolamine has also been reported, although thus far only in methanogenic Archaea from the genus *Methanococcoides* (77). The enzyme catalysing this reaction is unknown but could reasonably be assumed to be a methyltransferase within the MttB superfamily, which strain DMCF encodes in abundance. Further stepwise demethylation of dimethylethanolamine would yield ethanolamine, which can be transformed to acetaldehyde and ammonium within a bacterial microcompartment, also encoded in the strain DCMF genome. However, as all nitrogen in the provided choline was recovered within methylamine (127% ± 19% N recovery), this pathway seems less likely than transformation of choline to glycine betaine via betaine aldehyde in strain DCMF, as described in the main text.

*Demethylation of glycine betaine*

Glycine betaine (whether derived from choline or provided exogenously) is then likely demethylated, as has previously been reported in *Eubacterium limosum* (78), *Acetobacterium* spp. (79–81), and *Sporomusa* spp. (51). Demethylation is likely catalysed by a glycine betaine:corrinoid methyltransferase, encoded by a non-pyrrolysine member of the MttB superfamily (82). The glycine betaine:corrinoid methyltransferase ($MT_1$, MtgB) transfers a methyl group to a cognate corrinoid protein, and a methyl-tetrahydrofolate methyltransferase (MtgA, $MT_2$) then transfers the methyl group from the corrinoid protein to an accepting compound, tetrahydrofolate (82). In *S. ovata* strain An4, the same two mtgB genes were suggested to carry out demethylation of both glycine betaine and dimethylglycine, forming sarcosine (methylglycine) (52), whilst in *Acetobacterium woodii*, the protein appears to be specific to glycine betaine only, as there was no subsequent demethylation from dimethylglycine to sarcosine (83). MtgB homologs in the strain DCMF genome were previously described in a genome-based metabolic model, suggesting the organism may be capable of growth with glycine betaine and dimethylglycine (24), and an amended list of candidate proteins is included in Table S5, based on homology to known glycine betaine methyltransferases. Whilst a dimethylglycine methyltransferase has not yet been conclusively described in the literature, the alternative dimethylglycine dehydrogenase enzyme could not be identified in the strain DCMF genome, lending support to the suggestion by Visser *et al* (52) of catalysis by a methyltransferase.

*Reductive cleavage of sarcosine*

The genetic potential for reductive cleavage of glycine betaine and sarcosine was also reported in this metabolic model (24). Presuming demethylation of glycine betaine to sarcosine, DFE cultures amended with sarcosine + $H_2$ were set up to help verify this metabolite as a pathway intermediate of choline and glycine betaine catabolism, as it could not be observed at any stage of growth. The production of methylamine, acetate, and strain DCMF cells was consistent the proposed pathway. The apparent ability of strain DCMF to utilise $H_2$ as an electron donor for reductive cleavage of sarcosine was at odds with its inability to grow with the classic acetogenic substrates $H_2$ + $CO_2$. The genome does encode a putative membrane-bound NiFe hydrogen uptake hydrogenase (HyaABCD, Ga0180325_111497-9, Ga0180325_111503) which may be utilised to provide reducing equivalents for the sarcosine reductase.

Given the presence of putative glycine betaine reductases in the genome (Ga0180325_115251 and Ga0180325_115252s54) (24), DFE cultures were then amended with glycine betaine and $H_2$ to test whether reductive cleavage of glycine betaine was also possible. However, trimethylamine was not produced, despite $H_2$ depletion (Fig 2D). It is not yet clear which organisms in culture

DFE were utilising the $H_2$, as controls amended only with $H_2$ (i.e. glycine betaine-free or sarcosine-free) demonstrated no growth or acetogenesis. Concurrently, there was slightly higher acetate production observed in the glycine betaine + $H_2$ cultures (15 ± 0.6 mM, Fig 2D), compared to the $H_2$-free glycine betaine cultures (11 ± 0.4 mM; Fig 2C). It may be possible that $CO_2$ reduction by strain DCMF is enabled in the presence of glycine betaine and/or sarcosine, once other metabolic components (i.e. the WLP) are in use.

*Theoretical energy balance for quaternary amine metabolism*

Product formation and strain DCMF cell yields from the growth experiments with choline and glycine betaine were drawn together with the genomic information to generate a theoretical energy balance for consumption of choline and glycine betaine. The oxidation of two methyl groups from glycine betaine would yield 12 electrons (Eq. 6), of which two can be directed to reductive cleavage of sarcosine to yield one acetate and methylamine (Eq. 7). Given that sarcosine was not observed at any stage of growth, it is presumably rapidly cleaved. As eight electrons are required for acetate synthesis from bicarbonate (Eq. 8), the remaining 10 electrons equate to 1.25 acetate equivalents via bicarbonate reduction (Eq. 9), totalling 2.25 mol acetate equivalents and 1 mol methylamine per mole glycine betaine (Eq. 10). This approximately accords with the observed acetate (2.3 ± 0.1 mM per mole glycine betaine) and methylamine (0.9 ± 0.1 mM per mole glycine betaine) concentrations in glycine betaine-amended cultures.

The methylamine yield in choline-amended cultures (1.3 ± 0.2 mM per mole choline utilised), was also close to the theoretical yield based on the above equations. The metabolism of choline into glycine betaine liberates four electrons (Eq. 11), which equate to an additional 0.625 mol acetate for each mol of choline metabolised to glycine betaine (Eq. 12). Combining the choline to glycine betaine, and glycine betaine to acetate and methylamine equations results in a theoretical yield of 2.75 mol acetate equivalents per mole choline (Eq. 13), which is within one standard deviation of the observed 3.1 ± 0.4 mM acetate per mole choline utilised.

$(CH_3)_3N^+CH_2COO^- + 6\ H_2O \rightarrow (CH_3)NH_2^+CH_2COO^- + 2\ HCO_3^- + 14\ H^+ + 12\ e^-$ Eq. 6

$(CH_3)NH_2^+CH_2COO^- + 2\ H^+ + 2\ e^- \rightarrow (CH_3)NH_3^+ + CH_3COO^-$ Eq. 7

$2\ HCO_3^- + 9\ H^+ + 8\ e^- \rightarrow CH_3COO^- + 4\ H_2O$ Eq. 8

$2.5\ HCO_3^- + 11.25\ H^+ + 10\ e^- \rightarrow 1.25\ CH_3COO^- + 5\ H_2O$ Eq. 9

$(CH_3)_3N^+CH_2COO^- + H_2O + 0.5\ HCO_3^- \rightarrow 2.25\ CH_3COO^- + (CH_3)NH_3^+ + 0.75\ H^+$ Eq. 10

$(CH_3)_3N^+CH_2CH_2OH + H_2O \rightarrow (CH_3)_3N^+CH_2COO^- + 5\ H^+ + 4\ e^-$ Eq. 11

$HCO_3^- + 4.5\ H^+ + 4\ e^- \rightarrow 0.5\ CH_3COO^- + 2\ H_2O$ Eq. 12

$(CH_3)_3N^+CH_2CH_2OH + 1.5\ HCO_3^- \rightarrow CH_3NH_3^+ + 2.75\ CH_3COO^- + 1.25\ H^+$ Eq. 13

*Metabolism of methanol by strain DCMF*

Methanol catabolism in strain DCMF is proposed to be carried out via a methanol methyltransferase system, which is similar to the three-component system described above for glycine betaine, comprising a methanol:corrinoid methyltransferase (MtaB, $MT_1$), methyl-tetrahydrofolate methyltransferase (MtaA, $MT_2$), and cognate corrinoid protein (MtaC). While such methanol methyltransferase systems are relatively well-described in methanogenic archaea (84–91), there are only a few reports from acetogenic bacteria, namely in *Moorella thermoacetica* (92), *S. ovata* (52) and *A. woodii* (93). The strain DCMF genome encodes a number of methanol-specific methyltransferases and associated corrinoid proteins (Table S5). Within the strain DCMF genome, the closest homolog to MtaB from *S. ovata* and *A. woodii* is a methanol:corrinoid methyltransferase (Ga0180325_112644). It resides in a cluster containing a methanol-specific $MT_2$ homolog (Ga0180325_112641) and a MtbC homolog (Ga0180325_112642 and Ga0180325_112645).

As in both *S. ovata* and *A. woodii*, the putative $MT_2$ gene in strain DCMF is a methyl-tetrahydrofolate methyltransferase, rather than the MtaA methanol methyltransferase found in methanogens (52,93). In *A. woodii*, the resulting methyl-tetrahydrofolate is then be transformed via the WLP (93) and strain DCMF is expected to follow a similar metabolic route, leading to the formation of acetate as the sole product.

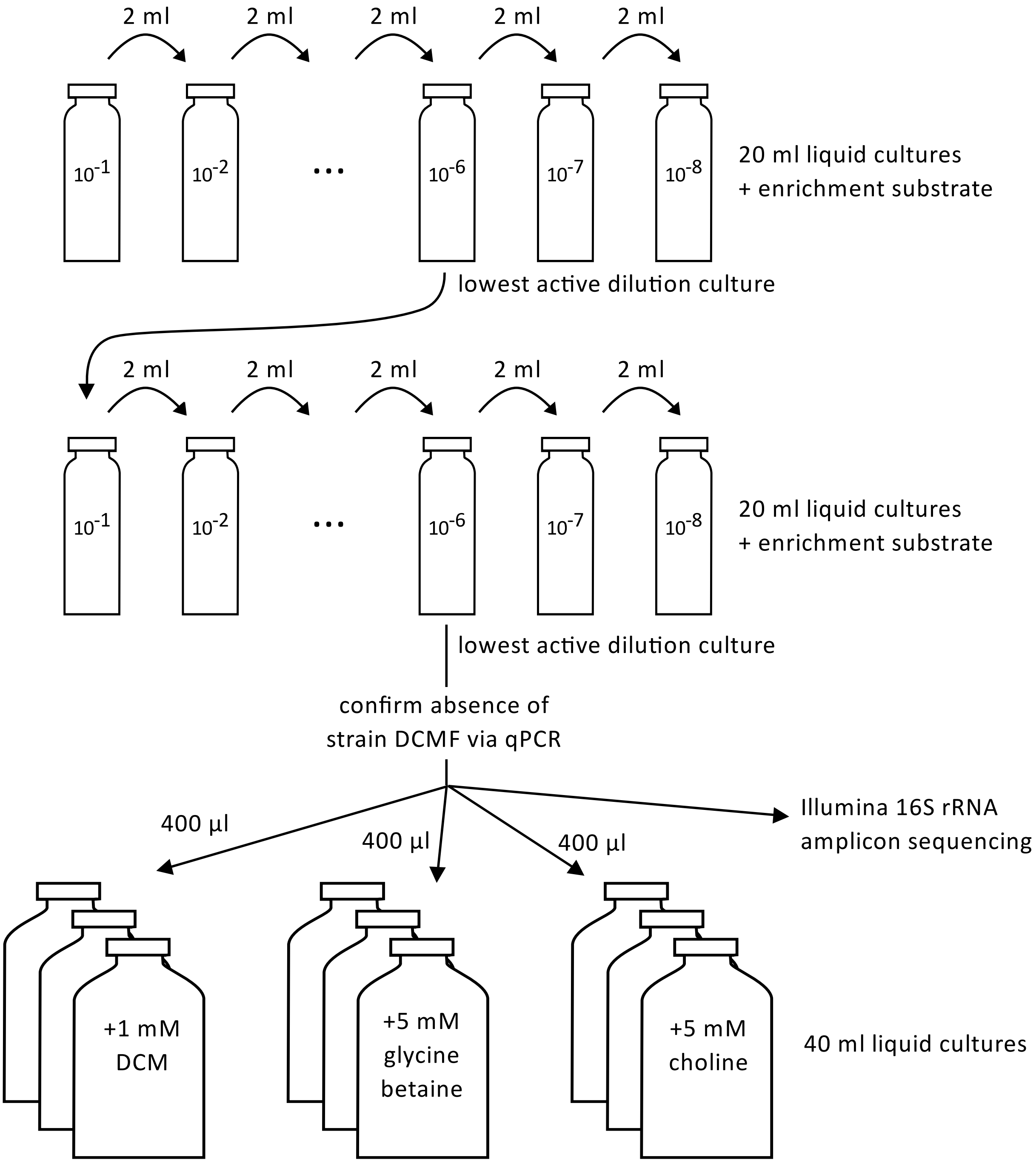


**Supplementary Figure 1. Overview of the exclusion cultivation method used to generate strain DCMF-free cohabitant enrichment cultures.**

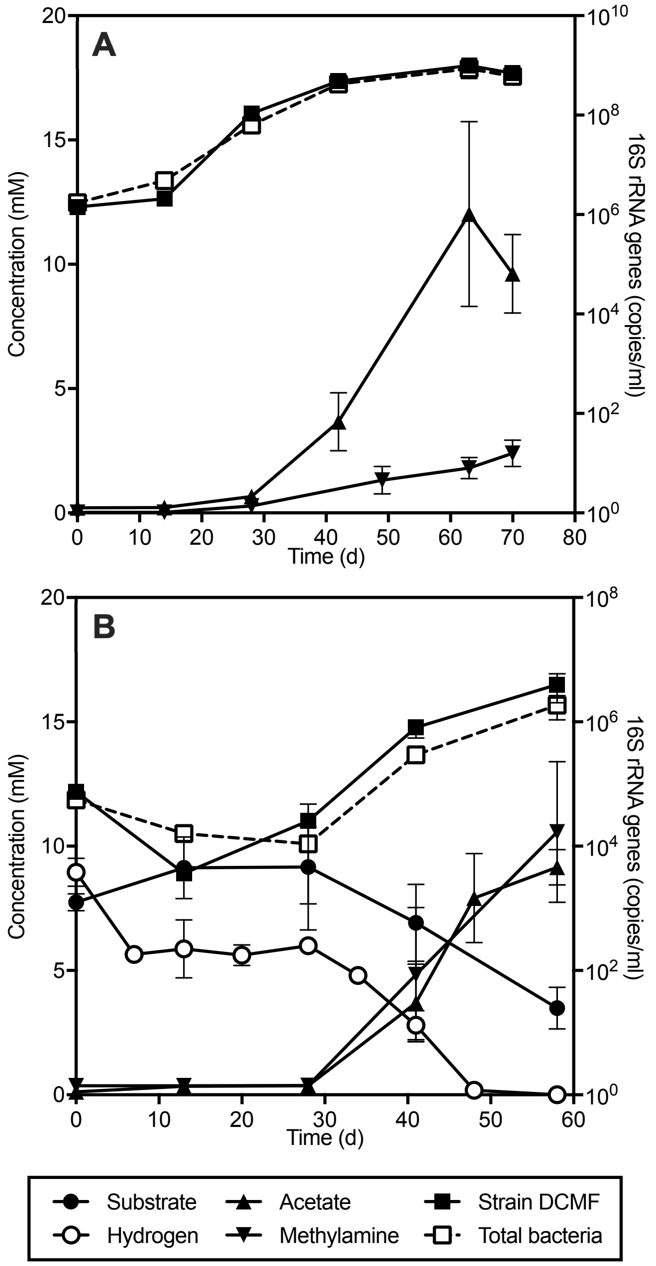


**Supplementary Figure 2. Strain DCMF is able to utilise (A) dimethylglycine and (B) sarcosine (methylglycine) + $H_2$ for growth.** Dimethylglycine could not be quantified. Strain DCMF growth was concomitant with an increase in acetate and methylamine. Substrate and product concentrations are quantified on the left y-axis; strain DCMF and total bacterial 16S rRNA gene copies are quantified on the right y-axis. Error bars represent standard deviation, *n* = 3.

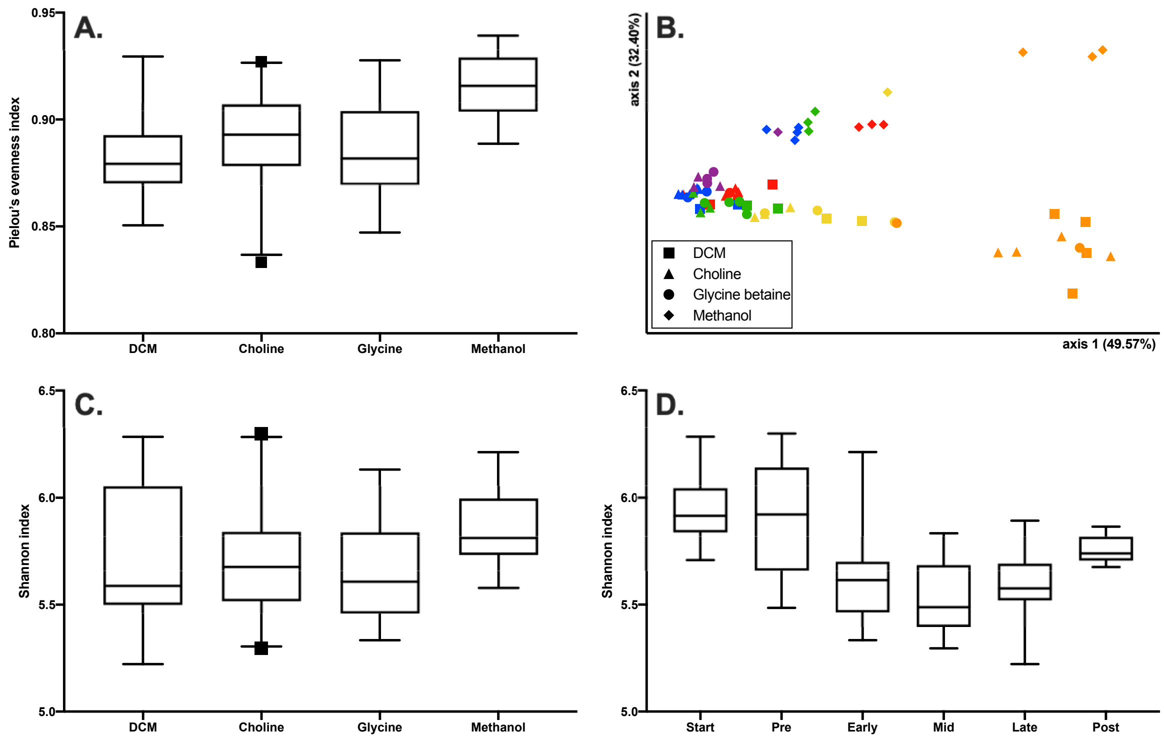


**Supplementary Figure 3. Shifts in the culture DFE community are driven by the stage of substrate consumption more than difference in substrate.** (A) Methanol-amended cultures had a significantly higher degree of evenness (adjusted p-value <0.01 in pairwise Kruskal-Wallis analysis of methanol compared to all other substrates) compared to cultures on the other three substrates, reflective of the lower relative abundance of strain DCMF in these cultures. (B) Principle components analysis plot of the weighted Unifrac distance matrix. Samples tended to group together based on substrate consumption proportion (colours) rather than differing substrate (shapes), although the methanol-amended community showed a higher degree of difference overall. Clusters emerged when samples were grouped by substrate consumption: inoculum/day 0 samples (red), pre (orange), early (yellow), mid (green), late (blue), or post (purple) (Table S3). There was no significant difference in the Shannon diversity index between samples grouped by substrate (C), but significant differences between samples when grouped by substrate consumption (D).

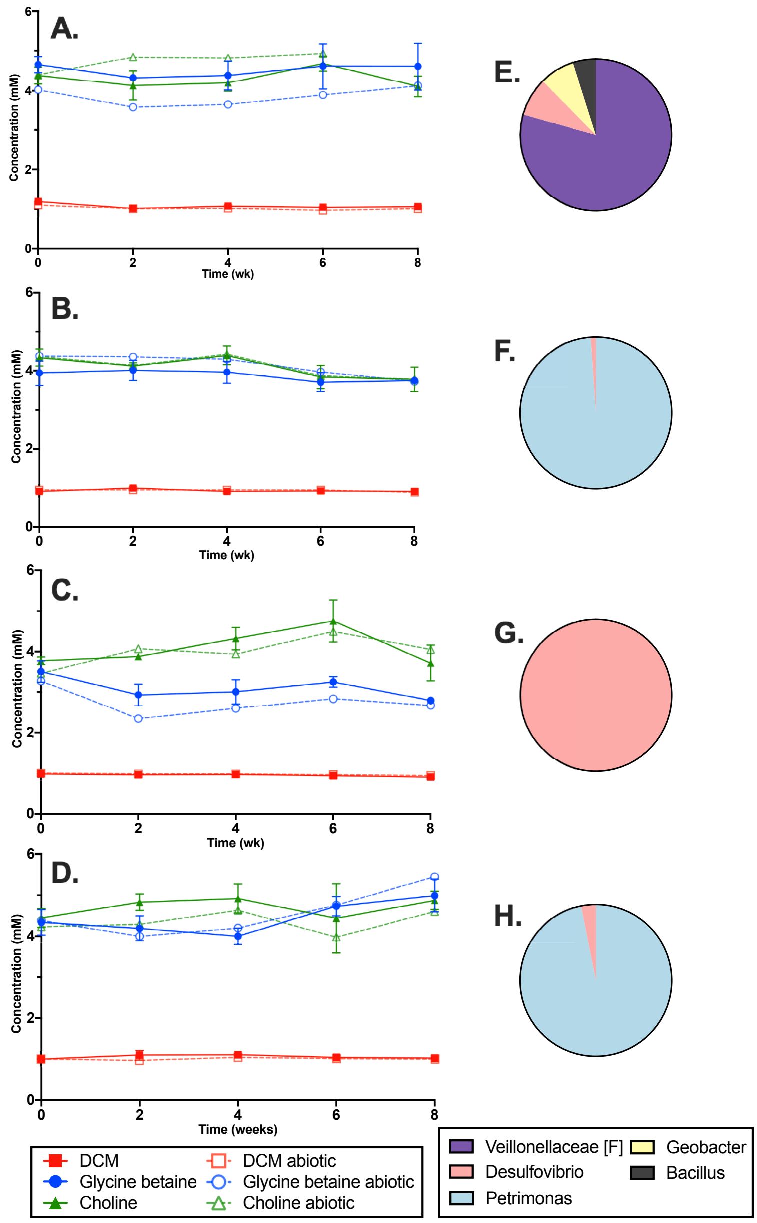


**Supplementary Figure 4, DFE community enrichments excluding strain DCMF cannot consume DCM, glycine betaine, or choline.** DFE cohabitants were enriched on casamino acids (A), glucose (B), peptone (C), yeast extract (D) to the exclusion of strain DCMF, then transferred back into medium containing the typical strain DCMF substrates (DCM, glycine betaine, choline). Active cultures $n$ = 3; abiotic $n$ = 1; error bars represent standard deviation. Community profiling was carried out via Illumina 16S rRNA gene amplicon sequencing with the lowest active dilution culture in the second round of dilution-to-extinction on casamino acids (E), glucose (F), peptone (G), yeast extract (H); i.e., the cultures used as inoculum for the cultures depicted in (A) to (D), respectively.

### Supplementary Tables

See separate Excel document containing Supplementary Tables 1 – 6.

### Supplementary Information References

60. Chaumeil P-A, Mussig AJ, Hugenholtz P, Parks DH. GTDB-Tk: a toolkit to classify genomes with the Genome Taxonomy Database. Bioinformatics. 2019;btz848.

68. Tan B, Jane Fowler S, Laban NA, Dong X, Sensen CW, Foght J, et al. Comparative analysis of

metagenomes from three methanogenic hydrocarbon-degrading enrichment cultures with 41 environmental samples. ISME J. 2015;9(9):2028–45.

69. Marchandin H, Jumas-Bilak E. The Family Veillonellaceae. In: Rosenberg E, Delong EF, Thompson F, editors. The Prokaryotes: Firmicutes and Tenericutes. Berlin: Springer-Verlag; 2014.

84. van der Meijden P, Jansen LPJM, Drift C, Vogels GD. Involvement of corrinoids in the

methylation of coenzyme M (2-mercaptoethanesulfonic acid) by methanol and enzymes from *Methanosarcina barkeri*. FEMS Microbiol Lett. 1983;19(2–3):247–51.
